# The molecular foundation of proprioceptor muscle-type identity

**DOI:** 10.1101/2022.07.29.501977

**Authors:** Stephan Dietrich, Carlos Company, Kun Song, Elijah David Lowenstein, Levin Riedel, Carmen Birchmeier, Gaetano Gargiulo, Niccolò Zampieri

## Abstract

The precise execution of coordinated movements depends on proprioception, the sense of body position in space. However, the molecular underpinnings of proprioceptive neuron subtype identities are not clear yet. In this study, we searched for molecular correlates of proprioceptor subtypes defined according to the identity of the muscle they innervate. We identified and validated signatures for subtypes monitoring the activity of back, abdominal, and hindlimb muscles. We found that proprioceptor muscle identity is acquired early in development and comprise programs controlling wiring specificity. Altogether this work paves the way for defining the mechanisms underlying the development of proprioceptor subtypes to the single muscle level and dissect their contributions to motor control.

## Introduction

Proprioception, the sense of body position in space, is critical for the generation of coordinated movements and reflexive actions. The primary source of proprioceptive information is represented by sensory neurons in the dorsal root ganglia (DRG), whose afferents innervate specialized mechanoreceptive organs detecting muscle stretch and tension (Proske and Gandevia, 2012). Proprioceptive sensory neurons can be anatomically and functionally divided on the basis of the identity of the muscle and receptor organ they innervate. First, during early development, proprioceptors innervate muscles and in order to precisely adjust motor output according to the biomechanical properties of their targets wire with neural circuits in the central nervous system (CNS) with exquisite specificity (Balaskas et al., 2020; Meltzer et al., 2021). In addition, at a receptor level, proprioceptors can be further distinguished into three subtypes - Ia, Ib, and II - by their selective contribution to either muscle spindles (MS; Ia and II) or Golgi tendon organs (GTO; Ib) (Zampieri and de Nooij, 2021). Most notably, Ia sensory afferents make monosynaptic connections to motor neurons controlling the activity of the same muscle, as well as synergist muscle groups, while avoiding motor neurons controlling the function of antagonist muscles, thus providing the anatomical substrate for the stretch reflex (Eccles et al., 1957; Mears and Frank, 1997). These precise patterns of connectivity are conserved in all limbed vertebrates and their assembly precedes the emergence of neural activity (Mendelsohn et al., 2015; Mendelson and Frank, 1991), implying that proprioceptive neurons are endowed from early developmental stages with molecular programs controlling critical features of their muscle-type identity, such as central and peripheral target specificity (Poliak et al., 2016; Shin et al., 2020). However, these determinants are still largely unknown, thus hindering efforts to define the mechanisms underlying the development of proprioceptive sensory neuron subtypes, the wiring of spinal sensorimotor circuits, and the contribution of muscle-specific proprioceptive feedback to motor control.

Single-cell transcriptomic efforts have revealed remarkable diversity among the major types of somatosensory neurons, while proprioceptors, despite their evident functional heterogeneity, seemed to represent a relatively more homogenous population (Chiu et al., 2014; Sharma et al., 2020; Usoskin et al., 2015). Recent studies aimed at characterizing the molecular nature of group Ia, Ib, and II neurons have revealed that signatures for receptor subtypes emerge late during development and are consolidated at postnatal stages (Oliver et al., 2021; Wu et al., 2021). However, the molecular basis of proprioceptor muscle-type identity remains elusive and so far only few markers for muscles in the distal hindlimb compartment have been identified (Poliak et al., 2016).

In this study, we used a single-cell transcriptomic approach that takes advantage of the somatotopic organization of proprioceptor muscle innervation to reveal the molecular profiles of cardinal muscle identities - epaxial and hypaxial - defined by peripheral connectivity to back and abdominal muscle groups at thoracic level, and lower back and hindlimb muscles at lumbar level. Our data show that muscle-type identity is acquired and consolidated during embryonic development and precedes the emergence of receptor character. In addition, we found that the identified molecular signatures comprise programs controlling defining features of proprioceptor muscle character, such as the specificity of muscle connectivity. In particular, differential expression of axon guidance molecules of the ephrin-A/EphA family discriminates epaxial and hypaxial muscle identities and elimination of ephrin-A5 function erodes the specificity of peripheral connectivity. Altogether, this study reveals that muscle-type identity is a fundamental aspect of proprioceptor subtype differentiation that is acquired during early development and includes molecular programs involved in the control of muscle target specificity.

## Results

### Transcriptome analysis at e15.5 reveals distinct proprioceptive clusters

In order to identify molecular correlates of proprioceptive sensory neurons (pSN) muscle identity we used transcriptome analysis of neurons isolated from thoracic and lumbar DRG at embryonic day (e) 15.5 At this stage proprioceptors have just reached muscle targets in the periphery and their central afferents are progressing toward synaptic partners in the ventral spinal cord (Extended Data Fig. 1a) (Zampieri and de Nooij, 2021). In addition, neurons collected from different segmental levels are predicted to reveal traits of epaxial and hypaxial pSN muscle identities, as the cell bodies of neurons innervating back and abdominal muscle groups are found in thoracic DRG, while the ones innervating lower back and hindlimb muscle groups in lumbar ones (Fig. 1a).

**Fig. 1.**
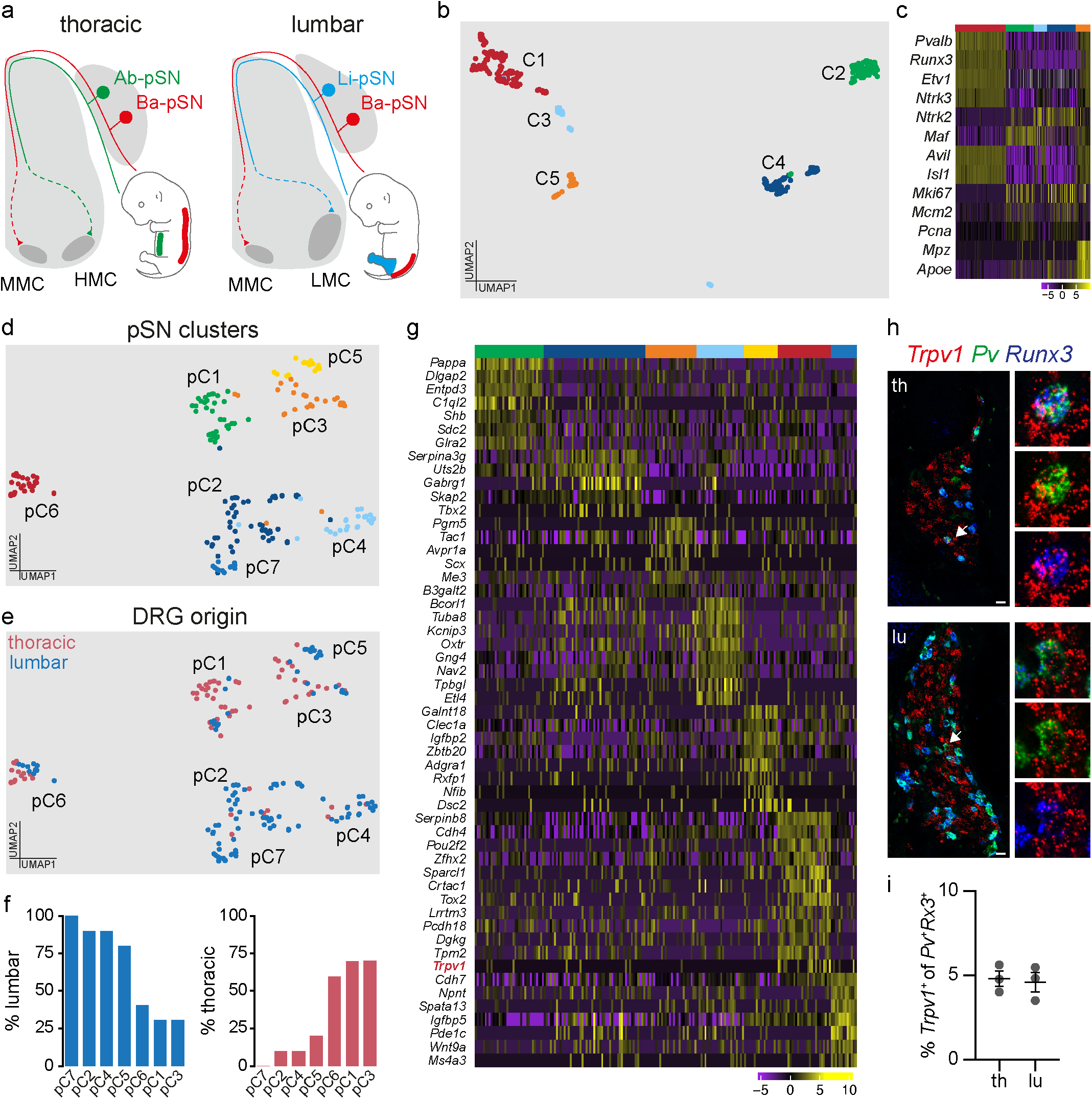
Single-cell transcriptome analysis of thoracic and lumbar proprioceptors at e15.5. **a)** Schematic illustrating central and peripheral connectivity of e15.5 proprioceptors at thoracic (left) and lumbar (right) spinal levels. Ab-pSN, abdominal muscles-connecting proprioceptors; Ba-pSN, back muscles-connecting proprioceptors; Li-pSN, hindlimb muscles-connecting proprioceptors; MMC, median motor column; HMC, hypaxial motor column; LMC, lateral motor column. **b)** UMAP visualization of tdTomato^+^ neuron clusters from *Pv^tdTom^* embryos at e15.5. **c)** Gene expression analysis (logcounts) of proprioceptors (*Pv*, *Runx3*, *Etv1*, *Ntrk3*), mechanoreceptors (*Ntrk2*, *Maf)*, postmitotic neurons (*Avil*, *Isl1*), proliferating neurons (*Mki67*, *Mcm2*, *Pcna*) and glial (*Mpz*, *Apoe*) markers. **d)** UMAP visualization of proprioceptor clusters identified from analysis of C1. **e)** UMAP visualization of proprioceptor clusters color-coded according to the thoracic (red) and lumbar (blue) origin of the cells. **f)** Percentage of proprioceptors originating from lumbar (left) and thoracic (right) DRG in different proprioceptor clusters**. g)** Differential gene expression analysis (logcounts) in proprioceptive clusters (pC1, green; pC2, dark blue; pC3, orange; pC4, light blue; pC5, yellow; pC6, red; pC7; blue). **h)** Representative single molecule fluorescent *in situ* hybridization (smFISH) images of thoracic (top) and lumbar (bottom) e15.5 DRG sections showing proprioceptors (*Runx3^+^*; *Pv^+^*) expressing *Trpv1*. Scale bar: 25 µm. **i)** Percentage of proprioceptors (*Runx3^+^*; *Pv^+^*) expressing *Trpv1* in thoracic and lumbar DRG at e15.5 (mean ± SEM, n = 3).

We took advantage of parvalbumin expression in proprioceptors and a small subset of cutaneous mechanoreceptors (de Nooij et al., 2013) to isolate 960 neurons - 480 from thoracic (T) levels 1 to 12 and 480 from lumbar (L) levels 1 to 5 - via fluorescence-activated cell sorting after dissociation of DRG from a BAC mouse line expressing tdTomato under the control of the parvalbumin promoter (*Pv^tdTom^*) (Kaiser et al., 2016) and processed them using the CEL-Seq2 protocol (Extended Data Fig. 1b and 1c) (Hashimshony et al., 2016). 519 neurons passed quality controls (see methods for details) and were found distributed into five molecularly distinct clusters (Fig. 1b and Extended Data Fig. 1d-f). Transcriptome analysis indicated that neurons in cluster (C) 1 represent proprioceptors, as they express general markers of proprioceptive identity (*Pv, Runx3, Etv1,* and *Ntrk3*). C2-C4 neurons present a signature consistent with mechanoreceptor identity (*Maf^+^*and *Ntrk2^+^*), with C3 neurons consisting of a postmitotic subset (*Isl1^+^* and *Avil^+^*) while C2 and C4 are characterized by proliferation markers (*Mki67^+^, Mcm2^+^,* and *Pcna^+^)*. Finally, C5 represents neurons contaminated with glial transcripts (*Mp*z*^+^* and *Apoe^+^*; Fig. 1c and Extended Data Fig. 1g) (Lallemend and Ernfors, 2012).

Next, to highlight differences between proprioceptors we re-clustered C1 neurons and obtained seven subsets (pC1-pC7; Fig. 1d and Extended Data Fig. 1h). In order to test whether anatomical provenance could point to proprioceptor muscle identities, we assigned anatomical origin to each cell. We found that neurons in pC2, pC4, pC5, and pC7 mainly originated from lumbar DRG and therefore could represent pSN connected to hindlimb muscles or the small subset of back muscles found at lumbar levels (lower back and tail muscles), while pC1, pC3, and pC6 present significant contribution from thoracic levels, where proprioceptors innervating back and abdominal muscles are located (Fig. 1a, e, and f) (Brink and Pfaff, 1980). We confirmed thoracic and lumbar origin at a transcriptional level by evaluating expression of *Hoxc10,* a gene defining lumbar identity (Philippidou and Dasen, 2013), and found that it closely recapitulated the anatomical assignment (Extended Data Fig. 1i). Next, we performed differential gene expression analysis and revealed distinct molecular signatures for each of these clusters (Fig. 1g). Surprisingly, we found that *Trpv1* is selectively enriched in neurons found in pC6 (Extended Data Fig. 1j). Trpv1 is a well-known marker of nociceptive/thermosensitive neurons and therefore is not expected to be expressed in proprioceptors (Mishra et al., 2011). Nevertheless, we confirmed the presence of *Pv*^+^; *Runx*3^+^; *Trpv*1^+^ DRG sensory neurons in e15.5 embryos, representing at this stage ∼ 5% of all proprioceptors, both at thoracic and lumbar levels (Fig. 1h and i).

### Embryonic expression of Trpv1 defines a subset of proprioceptors connected to back muscles

Next, to verify whether *Trpv1* expression in embryonic proprioceptors marks a discrete neuronal subtype we took advantage of mouse lines driving expression of Cre and Flp recombinases under control of the Trpv1 (*Trpv1^Cre^*) (Cavanaugh et al., 2011) and parvalbumin (*Pv^Flp^*) (Madisen et al., 2010) promoters to label neurons with an intersectional tdTomato reporter allele (*Trpv1*; *Pv; tdT, Ai65*) (Madisen et al., 2015).

Anatomical analysis of postnatal day (p) 7 spinal cords, DRG, and muscles from *Trpv1*; *Pv; tdT* mice revealed a well-defined subset of sensory neurons. In the spinal cord, we found labeling of central afferents targeting and making vGluT1^+^ synaptic contacts with ChAT^+^ motor neurons in the median motor column (MMC), both at thoracic and lumbar levels, which are known to selectively innervate back and lower back/tail muscles (Fig. 2a, b, Extended Data Fig. 2b and Supplemental Videos 1 and 2). In contrast, limb-projecting motor neurons in the lateral motor column (LMC) or abdomen-projecting motor neurons in the hypaxial motor column (HMC) received little, if any, input from *Trpv1*; *Pv; tdT* axons (Fig. 2a, 2b and Supplemental Video 1) (Jessell, 2000). In agreement with selective central innervation of neurons in the MMC, which is the only motor neuron column present at all rostro-caudal spinal levels, we observed labeling of a subset of parvalbumin^+^ neurons in cervical, thoracic, and lumbar DRG (Fig. 2c-e, and Extended Data Fig. 2c). In the periphery, we found labeling of type Ia, Ib, and II receptors in back but not abdominal muscles (Fig. 2f and Extended Data Fig. 2a). Finally, in order to test the overall specificity of lineage tracing in *Trpv1*; *Pv; tdT* mice, we analyzed reporter expression in the brain. We did not find any labeling asides from axons projecting to the dorsal column nuclei of the brainstem that are known to receive direct innervation from proprioceptive sensory neurons (Extended Data Fig. 2e).

**Fig. 2.**
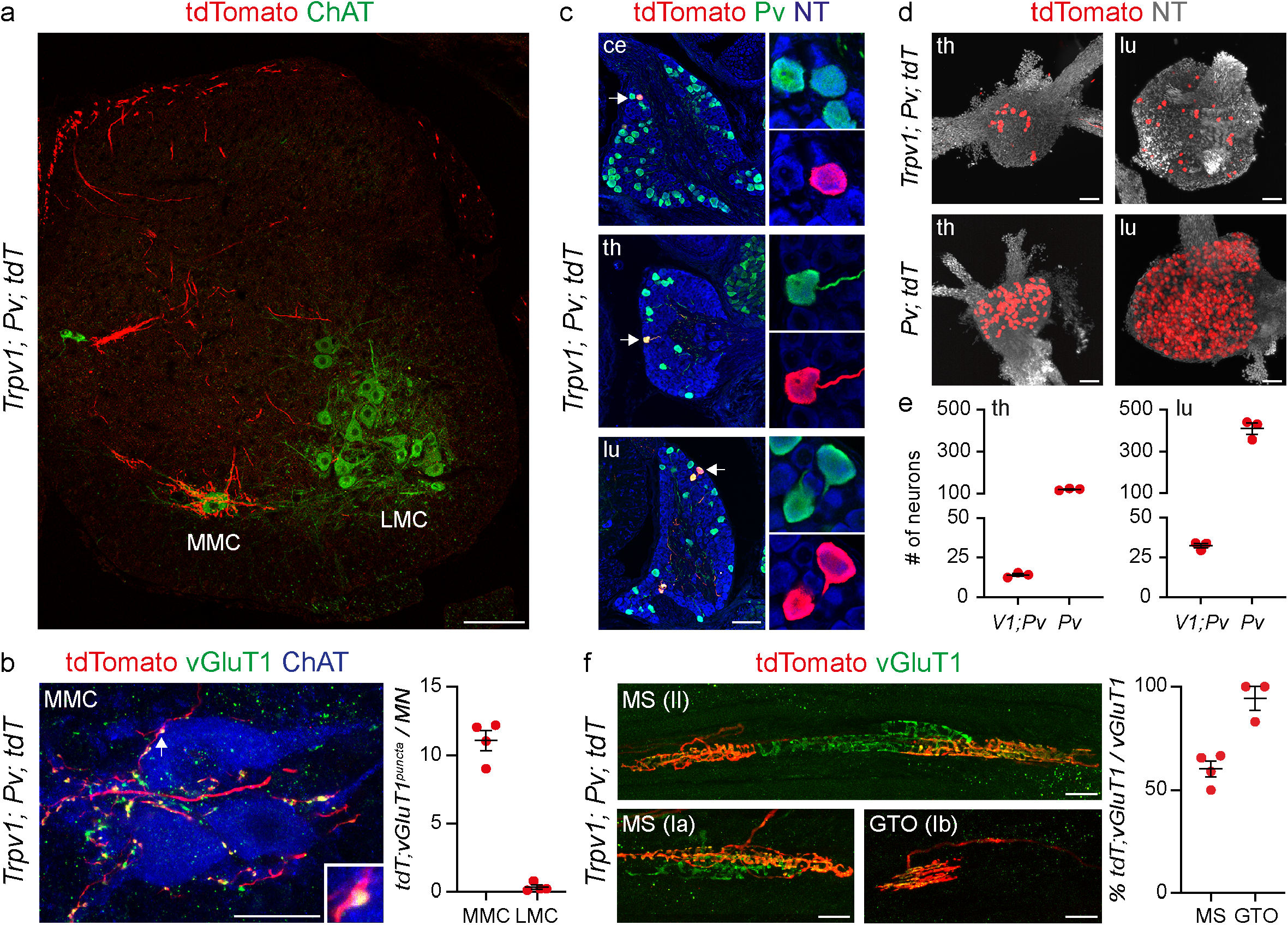
Genetic targeting of back muscles innervating proprioceptors. **a)** Representative image of tdTomato^+^ afferents in a lumbar spinal cord section from p7 *Trpv1^Cre^; Pv^Flp^; Ai65* mice. MMC, median motor column; LMC, lateral motor column. Scale bar: 100 µm. **b)** Representative image (MMC, left) and quantification (right) of tdTomato^+^; vGluT1^+^ presynaptic puncta juxtaposed to MMC or LMC neurons from p7 *Trpv1^Cre^; Pv^Flp^; Ai65* mice. Scale bar: 25 µm. **c)** Representative images of cervical, thoracic, and lumbar DRG sections showing tdTomato^+^; Pv^+^ neurons in p7 *Trpv1^Cre^; Pv^Flp^; Ai65* mice. Scale bar: 100 µm. **d)** Whole mount preparations of thoracic (left) and lumbar (right) DRG showing genetically labelled neurons from p1 *Trpv1^Cre^; Pv^Flp^; Ai65* (top) and *Pv^Cre^; Ai14* (bottom) mice. Scale bar: 100 µm. **e)** Number of tdTomato^+^ sensory neurons in DRG from p1 *Trpv1^Cre^; Pv^Flp^; Ai65* and *Pv^Cre^; Ai14* at thoracic (T1-T12, left) and lumbar (L1-L5, right) levels (n = 3, mean ± SEM). **f)** Representative images (left) and quantification (right) of tdTomato^+^ group Ia, II, and Ib afferents in muscle spindles (MS) and Golgi tendon organs (GTO) from the erector spinae muscle of *Trpv1^Cre^; Pv^Flp^; Ai65* mice. Scale bar: 25 µm.

In addition, we assessed whether lineage tracing from the Trpv1 promoter (*Trpv1; tdT. Ai14*) (Madisen et al., 2010) would also capture the same population of proprioceptors. Indeed, we observed labeling of a subset of Pv^+^ neurons in cervical, thoracic, and lumbar DRG, whose central afferents selectively targeted MMC neurons at all segmental levels (Extended Data Fig. 2f, g). Altogether these data show that Trpv1 expression in embryonic proprioceptors defines a subset of proprioceptive sensory neurons selectively innervating back muscles.

### Molecular signatures of proprioceptor muscle-type identities

The opportunity to genetically access a defined subset of proprioceptors defined by their connectivity to the back muscle compartment prompted us to further investigate the molecular identity of back- (Ba-pSN), abdominal- (Ab-pSN), and hindlimb-innervating (Li-pSN) neurons. To this end, we dissociated DRG and manually picked 576 tdTomato^+^ neurons from thoracic and lumbar levels of *Pv; tdT* (*Pv^Cre^*; *Ai14*; 96 thoracic and 96 lumbar neurons) (Hippenmeyer et al., 2007), and *Trpv1*; *Pv; tdT* mice (192 thoracic and 192 lumbar neurons) at p1 and performed single-cell transcriptome analysis (Fig. 3a and Extended Data Fig. 3a). 244 cells passed quality control criteria (see methods for details) and were found distributed into four clusters expressing high levels of general proprioceptive markers (Fig. 3b and Extended Data Fig. 3b-e). Cluster C1 presented signs of glia contamination and was excluded from subsequent analysis (Extended Data Fig. 3f). For the remaining clusters, we used mouse line and segmental level of origin of each neuron as means to assign a presumptive muscle identity. We found that the majority of cells picked from *Trpv1*; *Pv; tdT* mice, thus bona fide Ba-pSN, were found in C2 (Fig. 3c, d and Extended Data Fig. 3g). The majority of lumbar neurons from *Pv; tdT* mice, putative Li-pSN, were found in C3 and the remaining thoracic neurons, by exclusions putative Ab-pSN, in C4 (Fig. 3c, d, and Extended Data Fig. 3g).

**Fig. 3.**
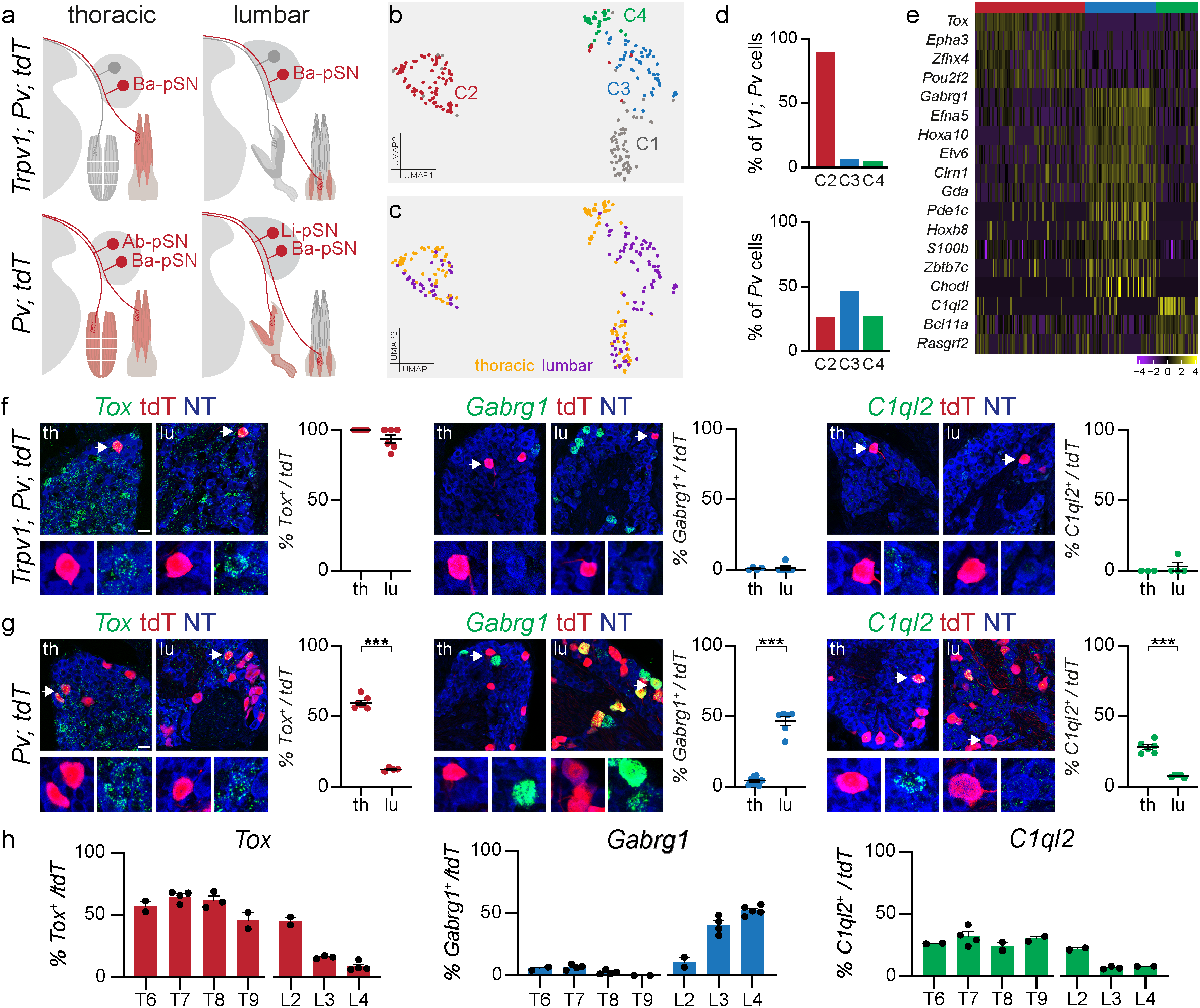
Molecular profiles of back-, adnominal- and hindlimb-innervating proprioceptors. **a)** Schematics illustrating labeling of the Ba-pSN subset captured in *Trpv1^Cre^; Pv^Flp^; Ai65* and all proprioceptors in *Pv^Cre^; Ai14* at thoracic and lumbar levels. **b)** UMAP visualization of cell clusters after transcriptome analysis of tdTomato^+^ DRG neurons from p1 *Trpv1^Cre^; Pv^Flp^; Ai65* and *Pv^Cre^; Ai14* mice at thoracic and lumbar levels. **c)** UMAP visualization of cell clusters color-coded according to the anatomical origin (thoracic, yellow; lumbar, purple) of neurons. **d)** Bar graph illustrating the percentage of *Trpv1^Cre^; Pv^Flp^; Ai65* (top) and *Pv^Cre^; Ai14* (bottom) cells found in clusters C2 (red), C3 (blue), C4 (green). **e)** Differential gene *e*xpression analysis (logcounts) for clusters C2 (red), C3 (blue), and C4 (green). **f)** Representative smFISH images and quantification of *Tox* (C2), *Gabrg1* (C3), and *C1ql2* (C4) expression in tdTomato^+^ thoracic and lumbar DRG neurons from p1 *Trpv1^Cre^; Pv^Flp^; Ai65* mice (each point represents one animal, mean ± SEM). Scale bar: 25 µm. **g)** Representative smFISH images and quantification of *Tox* (C2), *Gabrg1*(C3), and *C1ql2* (C4) expression in tdTomato^+^ thoracic and lumbar DRG neurons from p1 *Pv^Cre^; Ai14* mice (each point represents one animal, mean ± SEM, t-test, *** p < 0.001). Scale bar: 25 µm. **h)** Distribution of *Tox* (C2), *Gabrg1*(C3), and *C1ql2* (C4) in tdTomato^+^ neurons from thoracic and lumbar DRG of p1 *Pv^Cre^; Ai14* mice (each point represents one animal, mean ± SEM).

In addition, lumbar origin of each neuron was independently confirmed by analysis *Hoxc10* expression (Extended Data Fig. 3h).

Differential gene expression analysis revealed molecular signatures for presumptive Ba-pSN, Ab-pSN, and Li-pSN (Fig. 3e). To validate these findings, we first analyzed the expression of the top differentially expressed genes, *Tox* (C2, “Ba-pSN”), *Gabrg1* (C3, “Li-pSN”), and *C1ql2* (C4, “Ab-pSN”) (Extended Data Fig. 3i), in back-innervating proprioceptors labelled in *Trpv1*; *Pv; tdT* mice. In agreement with the predicted identity, *Tox* expression was found in nearly all tdTomato*^+^*neurons at thoracic and lumbar levels, while *Gabrg1* and *C1ql2* were not (Fig. 3f). Second, we examined expression and DRG distribution in the overall proprioceptive population labelled in *Pv; tdT* mice. At thoracic levels, where proprioceptors innervating back and abdominal muscle groups are located, we observed *Tox* expressed in ∼ 60% of tdTomato*^+^* neurons and *C1ql2* in ∼ 28%. At lumbar levels, where limb-innervating proprioceptors are predominant, we found *Gabrg1* in ∼ 46% of tdTomato*^+^* neurons and *Tox* in ∼ 10% (Fig. 3g). Altogether these data indicate that *Tox* and *C1ql2* are expressed within thoracic DRG and *Gabrg1* in lumbar DRG with frequencies expected for Ba-pSN, Ab-pSN and Li-pSN markers (Fig. 3h). Moreover, we found that expression of either *Gabrg1* or *Efna5*, another transcript differentially expressed in C3, covers ∼ 75% tdTomato*^+^* neurons at lumbar level, thus indicating that combination of multiple genes is necessary to define the hindlimb compartment (Extended Data Fig. 3j). Finally, in order to check whether effects of lineage tracing in *Pv; tdT* mice might influence the results, we analyzed expression of *Tox, Gabrg1,* and *C1ql2* in *Pv^+^* DRG neurons from wild-type mice and observed similar patterns and frequencies of expression at thoracic and lumbar levels (Extended Data Fig. 3k). Altogether, these data confirm that molecular markers of putative proprioceptor muscle subtypes identified with transcriptome analysis are expressed in thoracic and lumbar proprioceptive neurons from *Trpv1*; *Pv; tdT*, *Pv; tdT,* and wild-type mice with specificity and frequency consistent with back, abdominal and hindlimb muscle identities.

### Proprioceptor muscle identity emerges during early development

In order to further validate these observations and directly link molecular identity to muscle identity, we investigated expression of markers in proprioceptors subtypes identified by their muscle connectivity. To this end, we examined *Tox* (C2, “Ba-pSN”), *Gabrg1* (C3, “Li-pSN”), and *C1ql2* (C4, “Ab-pSN”) expression in DRG neurons retrogradely labeled after cholera toxin B (CTB) injection in representative back (erector spinae, ES) and hindlimb (gastrocnemius, GS; tibialis anterior, TA) muscles. We found that the majority of CTB^+^; *Pv^+^* neurons connected to ES expressed *Tox*, but neither *Gabrg1* nor *C1ql2* (Fig. 4a and Extended Data Fig. 4a). Conversely, proprioceptors labelled after CTB injections in hindlimb muscles expressed *Gabrg1*, but neither *Tox* nor *C1ql2* (Fig. 4b and Extended Data Fig. 4a). Altogether, these data show that genetic tracing and retrograde labeling experiments validated the findings of transcriptome analysis.

**Fig. 4.**
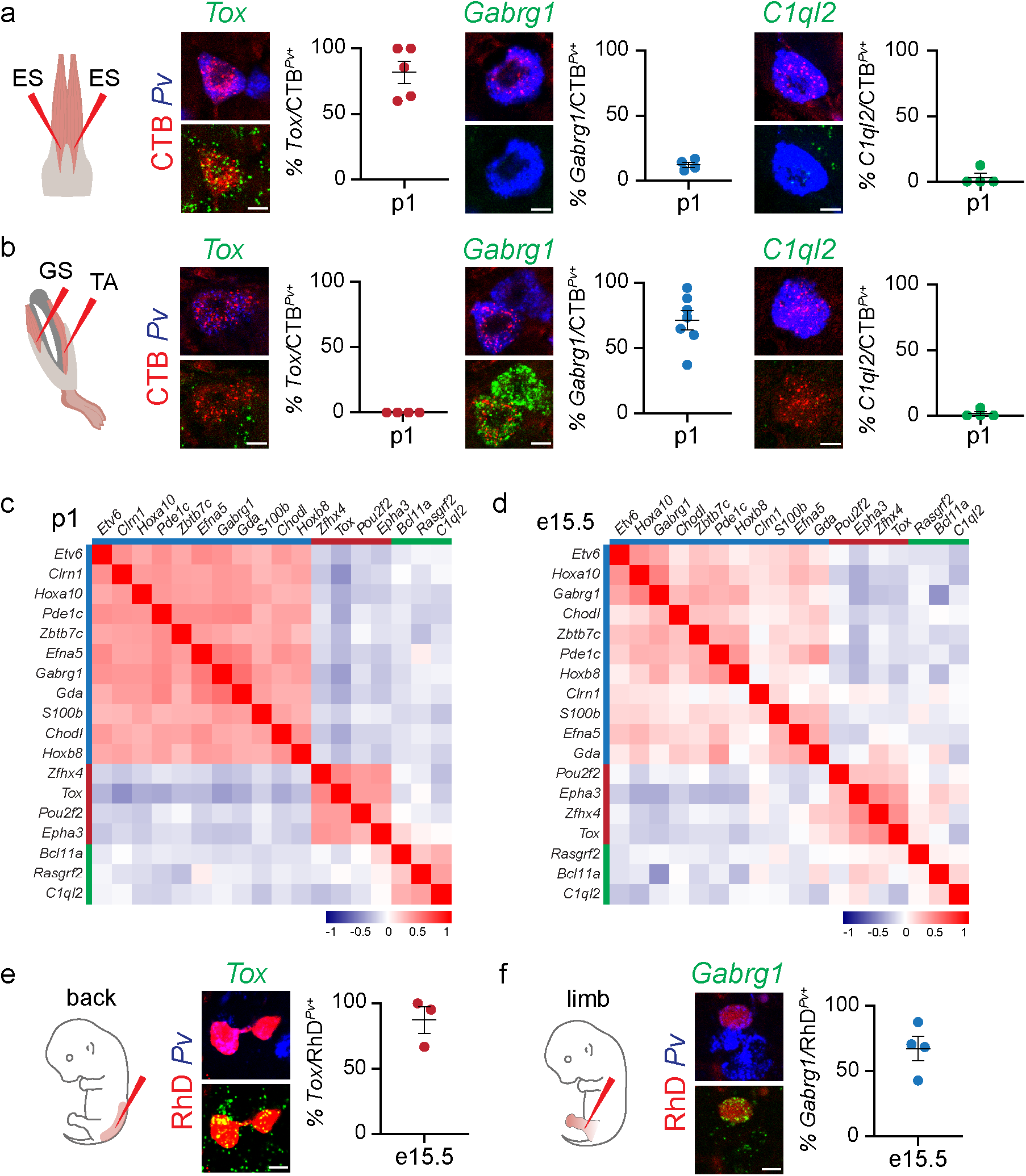
Proprioceptor muscle identity emerge at early developmental stages. **a and b)** Representative smFISH images and quantification of *Tox* (C2), *Gabrg1* (C3), and *C1ql2* (C4) expression in Pv^+^ sensory neurons retrogradely labelled after cholera-toxin B (CTB) injection in back (**a**; erector spinae, ES) and hindlimb (**b**; gastrocnemius, GS and tibialis anterior, TA) of p1 wild-type mice (each point represents one animal, mean ± SEM). Scale bar: 10 µm. **c and d)** Heatmaps representing pairwise gene expression correlation values for Ba-pSN (red), Ab-pSN (green), and Li-pSN (blue) molecular signatures at p1 (top) and e15.5 (bottom; Pearson’s r using logcounts). **e and f)** Representative smFISH images and quantification of *Tox* (C2), and *Gabrg1* (C3), expression in Pv^+^ sensory neurons retrogradely labelled after rhodamine-dextran (RhD) injection in e15.5 back (**e**) and hindlimb (**f**) muscles of wild-type mice (each point represents one animal, mean ± SEM). Scale bar: 10 µm.

Next, we asked whether gene expression profiles characterizing proprioceptor muscle identity at p1 were already present at earlier developmental stages. We analyzed correlation in expression of transcripts defining Ba-, Ab-, and Li-pSN identities at p1 and e15.5. As expected, we found high correlation at p1, in addition, strong co-expression patterns of the same signature genes are were also observed at e15.5 indicating that molecular features defining proprioceptor muscle identities are already present during embryonic development (Fig. 4c and d).

To confirm this finding, we examined expression of *Tox* and *Gabrg1,* in e15.5 DRG neurons retrogradely labeled after rhodamine-dextran (RhD) injection either in back or hindlimb muscles. As previously observed for postnatal stages, we found that expression of *Tox* and *Gabrg1* in embryonic proprioceptors is predictive of their specific peripheral connectivity patterns, with *Tox* labeling RhD^+^; *Pv^+^* back-innervating neurons and *Gabrg1* hindlimb-innervating ones (Fig. 4e, f and Extended Data Fig. 4b). Finally, we examined expression of p1 muscle-type markers (*Tox*, *Gabrg*, and *C1ql2*) in proprioceptor clusters identified at e15.5. We found that *Tox* expression characterizes three clusters (pC3, pC5 and pC6) of which two have predominant thoracic component, including the *Trpv1^+^* neurons in pC6, that represent Ba-pSN (Fig. 1d, e, f, g and Extended Data Fig. 1k). Consistent with Li-pSN, *Gabrg1* was found in two clusters (pC2 and pC7) whose neurons originate mainly from lumbar DRG, while *C1ql2* expression characterizes pC1 the only cluster formed by a majority of thoracic neurons, thus supporting Ab-pSN identity (Fig. 1d, e, f, g and Extended Data Fig. 1k).

Altogether these data indicate that molecular profiles of proprioceptor muscle subtypes identified at p1 are already present at e15.5 and part of developmental programs arising at embryonic stages before end-organ receptor identity consolidates (Oliver et al., 2021; Wu et al., 2021). Indeed, expression of molecular signatures recently identified for Ia, Ib and II receptor subtypes do not start being correlated in our datasets until p1 (Extended data Fig. 5).

### Ephrin-A/EphA signaling controls proprioceptor muscle targeting

The presence of molecular correlates of proprioceptor muscle character at early developmental stages suggests that signature genes defining different subtypes may be involved in the acquisition of their identities. Strikingly, the expression of *Efna5* and *Epha3* - members of the ephrin-A and EphA family of axon guidance ligands and receptors - distinguishes Ba- and Li-pSN (Fig. 3e, 4c, d and Extended Data Fig. 6a). Moreover, we found that other members of the EphA receptor family (*Epha4*, *Epha5*, and *Epha7*) are also differentially expressed in proprioceptor clusters, both at e15.5 and p1 (Fig. 5a). We validated these findings *in vivo* by characterizing expression of *Efna5* and *Epha3* in proprioceptors labelled in *Pv; tdT* and *Trpv1*; *Pv; tdT* mice at e15.5 and p1 (Fig. 5b and Extended Data Fig. 6b-d). In addition, we further confirmed these data by analyzing *Efna5* and *Epha3* expression in *Pv*^+^ sensory neurons retrogradely labelled after CTB injection in back and hindlimb muscles. We found that the majority of CTB^+^; *Pv^+^*neurons connected to hindlimb muscle expressed *Efna5* but not *EphA3* (Fig. 5c). Conversely, all the proprioceptors labelled after CTB injections in ES muscle expressed *Epha3*, but not *Efna5* (Fig. 5d). These data show that *Efna5* and *Epha3* are differentially expressed in Ba- and Li-pSN neurons, suggesting a function in controlling target specificity, an intriguing possibility considering the prominent role of ephrins and their receptors in axon guidance during development of the nervous system (Kania and Klein, 2016). First, in order to test whether ephrin-A5 controls proprioceptor peripheral connectivity, we injected CTB in hindlimb muscles of mice lacking ephrin-A5 function (*Efna5−/−*; Fig. 5e) (Frisén et al., 1998). First, to assess labeling specificity and whether elimination of ephrin-A5 was affecting motor neuron connectivity we examined the position and number of retrogradely labelled motor neurons. As previously reported, we did not find any significant difference in motor neuron muscle connectivity in *Efna5−/−* mice (Bonanomi et al., 2012) (Extended Data Fig. 6e-g).

**Fig. 5.**
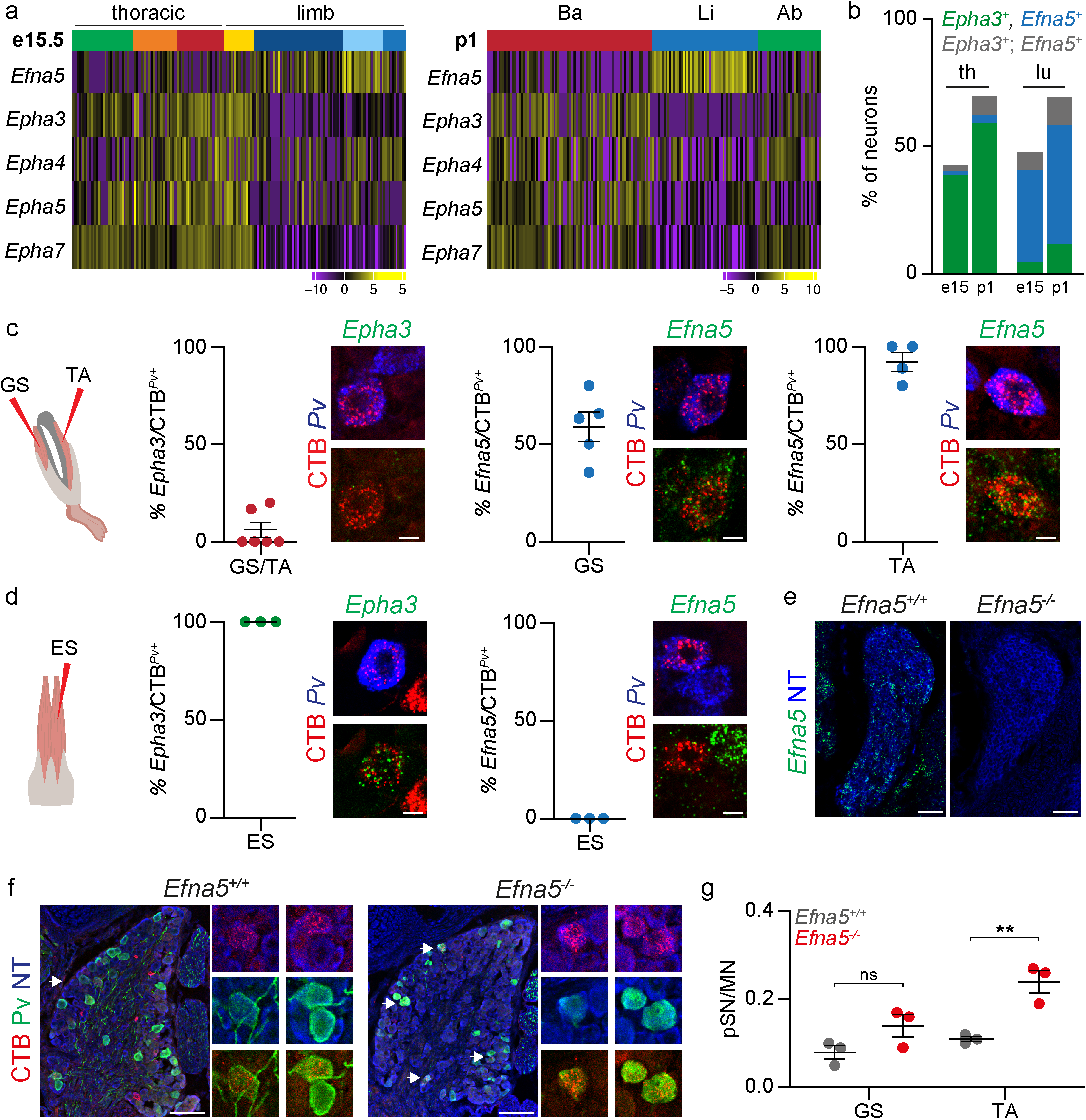
Elimination of ephrin-A5 function erodes the specificity of muscle connectivity. **a)** Gene expression analysis (logcounts) of ephrin-A/EphA family members differentially expressed in proprioceptor clusters identified at e15.5 (thoracic origin: green, orange, red; lumbar origin: dark blue, blue, light blue, and yellow) and p1 (Back: red; hindlimb: blue; abdominal green). **b)** Percentage of Pv^+^ or tdTomato^+^ sensory neurons expressing *Epha3* (green), *Efna5* (blue) or *Epha3; Efna5* (gray) at thoracic and lumbar levels in either e15.5 or p1 *Pv^Cre^; Ai14* mice. **c)** Representative smFISH images and quantification of *Epha3* (left) and *Efna5* (center/right) expression in Pv^+^ sensory neurons retrogradely labelled after CTB injection in gastrocnemius (GS), and tibialis anterior (TA) muscles of p1 wild-type mice (each data point represents one animal, mean ± SEM). Scale bar: 10 µm. **d)** Representative smFISH images and quantification of *Epha3* (left) and *Efna5* (right) expression in Pv^+^ sensory neurons retrogradely labelled after CTB injection in the erector spinae (ES) muscle of p1 wild-type mice (each point represents one animal, mean ± SEM). Scale bar: 10 µm. **e)** Representative smFISH images of *Efna5* expression in lumbar DRG of p1 *Efna5^+/+^* and *Efna5^−/−^* mice. Scale bar: 25 µm. **f)** Representative images of Pv^+^; CTB^+^ sensory neurons retrogradely labelled after CTB injection in the tibialis anterior (TA) muscle of p1 *Efna5^+/+^* (left) and *Efna5^−/−^* (right) mice. Scale bar: 100 µm. **g)** Ratio of proprioceptor (Pv^+^) per motor neuron labelled after CTB injection in the gastrocnemius (GS) and tibialis anterior (TA) muscles of p1 *Efna5^+/+^* (gray) and *Efna5^−/−^* (red) mice (each point represents one animal, mean ± SEM, t-test, ns p > 0.05, ** p < 0.01).

Next, we examined the number of retrogradely labeled proprioceptors, as well as the ratio of proprioceptor to motor neuron labeling, and found a significant increase in the number of neurons retrogradely labelled from the TA muscle, and a similar trend, although not significant, for the GS muscle, whose proprioceptors are only partially defined by *Efna5* expression (Fig. 5c, f, g and Extended Data Fig. 6g). Thus, these data show that elimination of ephrin-A5 function erodes the specificity of hindlimb muscle connectivity and indicate that the molecular signatures of muscle subtypes comprise programs controlling defining features of proprioceptor muscle-type identity.

## Discussion

This work defines the molecular signatures underlying proprioceptor subtypes defined by their muscle connectivity. We found that molecular distinctions emerge during embryonic development before the onset and consolidation of receptor character and comprise programs that control the specificity of muscle connectivity. These findings set the stage for defining the mechanisms controlling the acquisition of proprioceptor identity at a single muscle level and the generation of a new toolbox for analyzing the physiological roles of proprioceptor subtypes and define the contribution of sensory feedback from different muscle groups in the control of movement and the generation of the sense of body position in space.

We identified and validated molecular signatures for proprioceptor innervating cardinal muscle groups: back (*Tox*, *Epha3*), abdominal (*C1ql2*), and hindlimb (*Gabrg1, Efna5*). Markers for back and abdominal subtypes at thoracic level (*Tox* and *C1ql2*) account for almost the entire proprioceptor population in thoracic DRG (∼ 88%; Fig. 3g), thus indicating that our approach comprehensively captured most of the neurons innervating muscles at trunk level. In contrast, both *Gabrg1* (∼ 46%) and *Efna5* (∼ 66%) alone only capture about half of limb-innervating proprioceptors each, but together they account for about 75% of Pv^+^ sensory neurons in lumbar DRG (Fig. 3g, 5b, Extended Data Fig. 3j). The great anatomical complexity of limbs, comprising 39 different muscles in the mouse hindlimb (Charles et al., 2016), is consistent with a model requiring a combination of multiple molecules in order to represent the whole compartment. The presence of multiple clusters associated with general back and hindlimb identities at e15.5 indicate that their molecular makeup may already capture features of more fine-grained identities defined according to specific anatomical (i.e.: rostral vs. caudal back; dorsal vs. ventral limb) or functional (i.e.: synergist vs. antagonist) characteristics.

Upon acquisition of a generic proprioceptor fate (Marmigère and Ernfors, 2007), sensory neurons mature to develop functional features defined by their muscle and end-organ receptor identities. First, sensory axons navigate peripheral targets and innervate mechanoreceptive end-organs with precise ratios and distributions according to the biomechanical requirements of the innervated muscle (Banks et al., 2009). In addition, each proprioceptor subtype needs to establish specific sets of connections with multiple neural targets in the central nervous system in order to relay sensory feedback to motor circuits controlling the activity of relevant muscles (Chen et al., 2006; Mears and Frank, 1997). Our data support a model where proprioceptor muscle identity emerges as part of an embryonic genetic program controlling connectivity to its central and peripheral targets that is refined at later stages to include aspects of receptor-type character (MS and GTO), such as distinct physiological properties, whose diversification is influenced by neuronal activity (Wu et al., 2021). In support of this view, signatures of proprioceptor muscle-type identities are clearly evident from e15.5, while group Ia, Ib, and II molecular profiles have been shown to emerge later and consolidate during postnatal development (Oliver et al., 2021; Wu et al., 2019, 2021). Accordingly, molecular correlates defining receptor identity are not immediately evident in the muscle-type profiles we identified at e15.5, but start emerging at p1. A notable exception is represented by *Tox* and *Chodl,* which have been previously proposed to represent markers of two groups of type II afferents (II_2_ and II_4_) at early postnatal stages (Wu et al., 2021). These molecules define back (*Tox*) and hindlimb (*Chodl*) muscle subtypes in our analysis. Interestingly, groups II_2_ and II_4_ were found to be enriched in DRG at thoracic and lumbar levels respectively, thus confirming our results and indicating that the diversity observed in type II proprioceptors may already include signatures of muscle-type identity (Wu et al., 2021). Altogether, these observations suggest that “receptor” features become superimposed to “muscle” character already present since early development in order to generate the final functional subtype identity. Future studies building on these findings bear the promise to define the developmental processes controlling proprioceptor specification from general proprioceptive fate determination to the acquisition of muscle identity and maturation of physiological characteristics at receptor level.

The specificity with which proprioceptors innervate respective muscle targets in the periphery and synaptic partners in the central nervous system provides the circuit basis for the function of spinal sensorimotor circuits (Tuthill and Azim, 2018). Our data shows that the ephrin-A/EphA family of axon guidance molecules is an important regulator of proprioceptor peripheral connectivity. We found that differential expression of *ephrin-A5* and four EphA receptors (*EphA3*, *EphA4*, *EphA5*, and *EphA7*) delineate a distinction between hindlimb- and abdominal/back-projecting proprioceptors, and perturbation of ephrin-A5 function leads to an erosion in the specificity of muscle connectivity. The phenotype indicates that Efna5 may be part of a developmental program controlling the precision of muscle innervation. Ephrin signaling is known to have important roles in the guidance of somatosensory and motor axons to their peripheral targets (Kania and Klein, 2016). It has been shown that at early embryonic stages nascent sensory axons track along motor axons *en route* to their peripheral targets and trans-axonal interactions control navigation of sensory neurons to axial targets (Gallarda et al., 2008). In particular, interactions between EphA3/4 in motor axons and ephrin-A2/A5 in somatosensory axons have been shown to control innervation of the epaxial compartment by sensory neurons. In their absence, epaxial sensory nerves are re-routed to hypaxial targets (Wang et al., 2011). We observed a significant increase in the number of proprioceptors innervating the tibialis anterior muscle in mice lacking ephrin-A5, indicating that excessive limb muscle innervation might result from mistargeting of axons originally directed to another muscle whose identity remain elusive. However, we did not observe any difference in the connectivity to a representative back muscle, as the number of pSN retrograde labelled after ES injection in Efna5 −/− and control mice was unaffected (Extended Data Fig. 6h-k). Ephrin-A/EphA signaling also controls the choice of limb innervating motor neurons to invade either the dorsal or ventral half of the limb mesenchyme and could influence muscle by muscle dependence of proprioceptive axon innervation specificity (Bonanomi et al., 2012; Helmbacher et al., 2000; Kania and Jessell, 2003). Because of the intricacy of ephrin-Eph signaling (Kania and Klein, 2016), it will be necessary to carefully analyze the expression pattern and function of different ligands, receptors, and coreceptors in order to define the molecular logic governing guidance of proprioceptors to their specific muscle targets.

The importance of proprioceptive sensory feedback in motor control is clearly evident in mouse models where proprioceptor development or function is perturbed. Degeneration of muscle spindles in Egr3 mutant mice result in ataxia and, similarly, loss of most proprioceptors in absence of Runx3 function results in severe coordination phenotypes (Akay et al., 2014; Levanon et al., 2002; Tourtellotte and Milbrandt, 1998). Moreover, elimination of the mechanosensory transduction channel Piezo2 in proprioceptors leads to severely uncoordinated body movements and limb positions (Woo et al., 2015). Despite the critical role of proprioception for the generation of coordinated movement, it is still not understood how proprioceptive feedback from different muscles and receptor subtypes integrates with motor commands and other sources of sensory input to adjust motor output and generate the sense of body position in space (Pearson, 2004; Windhorst, 2007). This is mainly due to the fact that behavioral studies have been hampered by the lack of tools allowing precise access to different functional subtypes of proprioceptors. The identification of molecular signatures for proprioceptor muscle subtypes opens the way for the generation of new genetic and viral tools to selectively access distinct channels of proprioceptive information and bears the promise to determine their roles in motor control.

## Acknowledgements

We thank Liana Kosizki for technical support, Mathias Richter and the MDC Advanced Light Microscope facility for assistance with light sheet microscopy. We are grateful to Martyn Goulding and Mark Hoon for sharing mouse lines. We thank Susan Brenner-Morton for sharing antibodies. We thank Robert Manteufel, Ilka Duckert, and Florian Keim for animal care. We are grateful to Dario Bonanomi for helpful discussions; Nikos Balaskas, Joriene de Nooij, and members of the Zampieri laboratory for insightful comments on the manuscript. N.Z. was supported by DFG grant ZA 885/1-2; G.G. by Helmholtz (VH-NG-1153), KWF (NKI-2014-7208), and ERC (714922).

## Author contributions

Conceptualization, S.D. and N.Z.; Investigation, S.D., C.C., K.S., L.R., and E.D.L.; Formal analysis, S.D., C.C., L.R. and G.G.; Writing – Original Draft, S.D. and N.Z.; Writing – Review and Editing, S.D., C.C., K.S., L.R., E.D.L., C.B., G.G., and N.Z.; Supervision, C.B., G.G., and N.Z.

## Competing interests

The authors declare no competing interests.

## Data and materials availability

All unique reagents generated in this study are available from the Lead Contact without restriction. Datasets generated in the current study are deposited in GEO. The scripts and source data for the plots are available upon request.

## Methods

### Animal experimentation ethical approval

All experiments were performed in compliance with the German Animal Welfare Act and approved by the Regional Office for Health and Social Affairs Berlin (LAGeSo) under license numbers G0148/17 and G0191/18.

### Animal models

Mice were housed in standardized cages under 12h light-dark cycle with food and water *ad libitum*. For this study the following mouse lines were used *Pv^Cre^* (Hippenmeyer et al., 2007), *Pv^Flp^* (Madisen et al., 2015), *Pv^tdTom^*(Kaiser et al., 2016), *Trpv1^Cre-Basbaum^* (Cavanaugh et al., 2011), *Trpv1^Cre-Hoon^* (Mishra et al., 2011), *Ai14* (Madisen et al., 2010), *Ai65* (Madisen et al., 2015), and *Efna5−/−* (Frisén et al., 1998).

### Single-cell isolation

Dorsal root ganglia were dissected separately from thoracic (T1-T12) and lumbar (L1-l5) segments and collected in F12 medium with 10% FHS (Fetal horse serum) on ice. Next, DRGs were incubated in F12/FHS with 0,125% collagenase (Sigma C0130) for 1 hour (p1) or 30 min (e15.5) at 37°C. After 3 washes with PBS DRGs were transferred to 0,25% trypsin solution (Gibco 15050-065) and incubated for 15 min at 37°C. Afterwards, DRG were mechanically triturated using a fire polished Pasteur pipette until a homogenized solution was visible followed by a centrifugation step at 200 x g for 10 min. The final cell pellet was resuspended in HBSS (10 mM HEPES, 10 mM Glucose) and the resulting cell suspension either applied to fluorescence-activated cell sorting (FACS) (e15.5) or manual cell picking under an inverted fluorescent microscope (p1). Single tdTomato^+^ cells were sorted into individual wells containing lysis buffer and stored at −80°C until further processing.

### Single-cell RNA sequencing

For cDNA library preparation the CEL-Seq2 protocol was used as previously described (Hashimshony et al., 2016). We sequenced 960 cells (480 from T1-T12 and 480 from L1-L5) at e15.5 and 576 (96 thoracic and 96 lumbar from *Pv^Cre^; Ai14*; 96 thoracic and 96 lumbar from *Trpv1^Cre-Basbaum^; Pv^Flp^; Ai65*; 96 thoracic and 96 lumbar from *Trpv1^Cre-Hoon^; Pv^Flp^; Ai65*) at p1. The libraries were sequenced on an Illumina NextSeq500 platform with high-output flow cells by the Next Generation Sequencing Core Facility of the Max-Delbrück Center for Molecular Medicine.

### Single-cell analysis

For both data sets (e15.5 and p1) we used the scruff v1.4.0 package (R package version 1.12.0) to demultiplex, map, and generate count matrices. Then, we evaluated each data set statistics using Scater v1.14.6 R package. To increase the quality of the experiments, we individually removed low-quality cells based on low total gene counts (> quantile 0.3), low gene abundance (> quantile 0.3), and high mitochondrial gene values cells (< quantile 0.75). 519 out of 960 e15.5 cells and 244 out of 576 p1 cells passed quality control criteria. After log-normalization, we used the scran v1.14.1 buildKNNGraph and cluster_walktrap functions with default parameters to define each data-set cell populations and subclusters. Finally, we assigned gene markers to each population using findMarkers function from the scran with default parameters. For single cell analysis R v3.6.2 environment was used to generate the results, statistical analysis and graphical evaluation of the datasets.

### Dissection and tissue processing

Postnatal mice were anesthetized by intraperitoneal injection of 120 mg/kg ketamine and 10 mg/kg xylazine and transcardially perfused with PBS and 4% PFA in 0,1 M phosphate buffer. To expose the spinal cord a ventral laminectomy was performed and the tissue post-fixed O/N in 4% PFA at 4°C. The next day tissue was washed three times with ice-cold PBS and transferred to 30% sucrose in PBS for cryoprotection at 4°C O/N. Tissue was embedded in Tissue-Tek OCT embedding compound and stored at −80°C. 16 µm tissue sections for immunohistochemistry were acquired at a cryostat, dried for 1 hour and either directly used or frozen at −80°C.

### Immunohistochemistry and fluorescent in situ hybridization

For immunohistochemistry dry tissue sections were washed for 10 min with PBS followed by another 10 min incubation of 0.1% Triton-X-100 in PBS (0.1% PBX) for permeabilization. The following primary antibodies were diluted in 0.1% PBX and incubated O/N at 4°C: Ch-anti-Pv (1:5000, generous gift from Susan Brenner-Morton), Goat-anti-ChAT (1:200), GP-anti-vGluT1 (1:5000, generous gift from Susan Brenner-Morton), Rb-anti-dsRed (1:1000) and Rb-anti-RFP (1:500). Next, slides were washed three times for 5 minutes with 0.1% PBX followed by secondary antibody/NeuroTrace incubation for 1 hour at room temperature (RT). Secondary antibodies (Jackson Immuno Research Laboratories) and NeuroTrace (Life Technologies) were diluted in 0.1 % PBX as following: Cy3, Alexa488 (1:1000), Cy5 (1:250), and NeuroTrace (1:250). After staining with secondary antibodies slides were washed three times with 0.1% PBX and subsequently mounted with Vectashield antifade mounting medium. For fluorescent *in situ* hybridization the RNAscope Multiplex Fluorescent Kit v2 (ACDBio) with a modified manufactures protocol was used. Tissue sections were acquired as described above. Sections were dried, fixed with ice-cold 4% PFA in PBS for 15 min and dehydrated in a series of 50%, 70% and 100% ethanol for 5 min each. Afterwards, sections were treated with hydrogen peroxide solution for 15 min at RT to block endogenous peroxidase activity followed by another wash with 100% ethanol for 5 min. Next either Protease IV (postnatal tissue) or Protease III (embryonic tissue and sections from CTB tracing experiments) was applied for 30 min at RT. After three washes with PBS probes were applied and hybridization performed in a humified oven at 40°C for 2 hours. The following probes were used in this study: Mm-Epha3-C1, Mm-Tox-C1, Mm-C1ql2-C1, Mm-Efna5-C2, Mm-Trpv1-C2, Mm-Pvalb-C2, Mm-Pvalb-C3, Mm-Gabrg1-C3, and Mm-Runx3-C3. Following hybridization, amplification was performed using Amp1, Amp2 and Amp3 each for 30 min at 40°C. For detection each section was treated sequentially with channel specific HRP (HRP-C1, HRP-C1, HRP-C3) for 15 min, followed by TSA mediated fluorophore (Akoya Bioscience, Opal 520, Opal 570, and Opal 690) binding for 30 min and final HRP blocking for 15 min (all steps at 40°C). When necessary additional immunostaining was performed as described above. For quantification cell bodies (evaluated by Nissl staining) colocalizing with ≥5 puncta were counted positive.

### Tissue clearing and light-sheet microscopy

Mice were anesthetized and transcardially perfused as described above. Afterwards, spinal cord and/or DRG were extracted after ventral laminectomy and postfixed in 4% PFA for 2 days at 4°C. DRG were kept separately and embedded into 1% low melt agarose in OptiPrep (Sigma) after post fixation. Tissue clearing was performed as previously described with modifications (Susaki et al., 2015). In short, tissue was transferred to CUBIC1 (25 wt% Urea, 25 wt% N,N,N’,N’-tetrakis(2-hydroxypropyl) ethylenediamine, 15 wt% Triton X-100) and incubated at 37°C shaking. Every other day CUBIC1 solution was exchanged until tissue appeared transparent (spinal cord ∼ 4 days, DRG ∼ 1-2 days). Afterwards, samples were washed for 1 day with PBS at RT, refractive index matched with EasyIndex (LifeCanvas Technologies) at 37°C and imaged with the ZEISS Light-sheet Z.1. For image analysis and video rendering Arivis Vision4D (Arivis AG) and Imaris (Oxford Instruments) was used.

### Retrograde labeling of proprioceptors and motor neurons

For retrograde labelling of p1 proprioceptors, mice were anesthetized with isoflurane and a small incision on the skin was made to expose the muscle of interest. 50 nl of a 1% solution of Alexa555-conjugated CTB (Life Technologies) was injected with a glass capillary into the desired muscles. Animals were sacrificed and perfused after 3 days. For retrograde labelling of e15.5 proprioceptors, embryos were dissected in ice-cold artificial cerebrospinal fluid and pinned down. Afterwards, skin from limb or back muscles was removed and 20% rhodamine dextran (Life Technologies) injected into the desired muscle using a pulled glass capillary. Afterwards, embryos were incubated in circulating oxygenated artificial cerebrospinal fluid (5% CO2, 95% O2) for 6 hours at 27°C and fixed with 4% PFA.

### Quantification and statistical analysis

Details for statistical analysis and number of samples are indicated in figure legends. Significance for t-tests was defined as * p < 0.05; ** p < 0.01; *** p < 0.001. Statistically analyses were performed using Prism - GraphPad v9.2.

---

**Extended Data Fig. 1.**
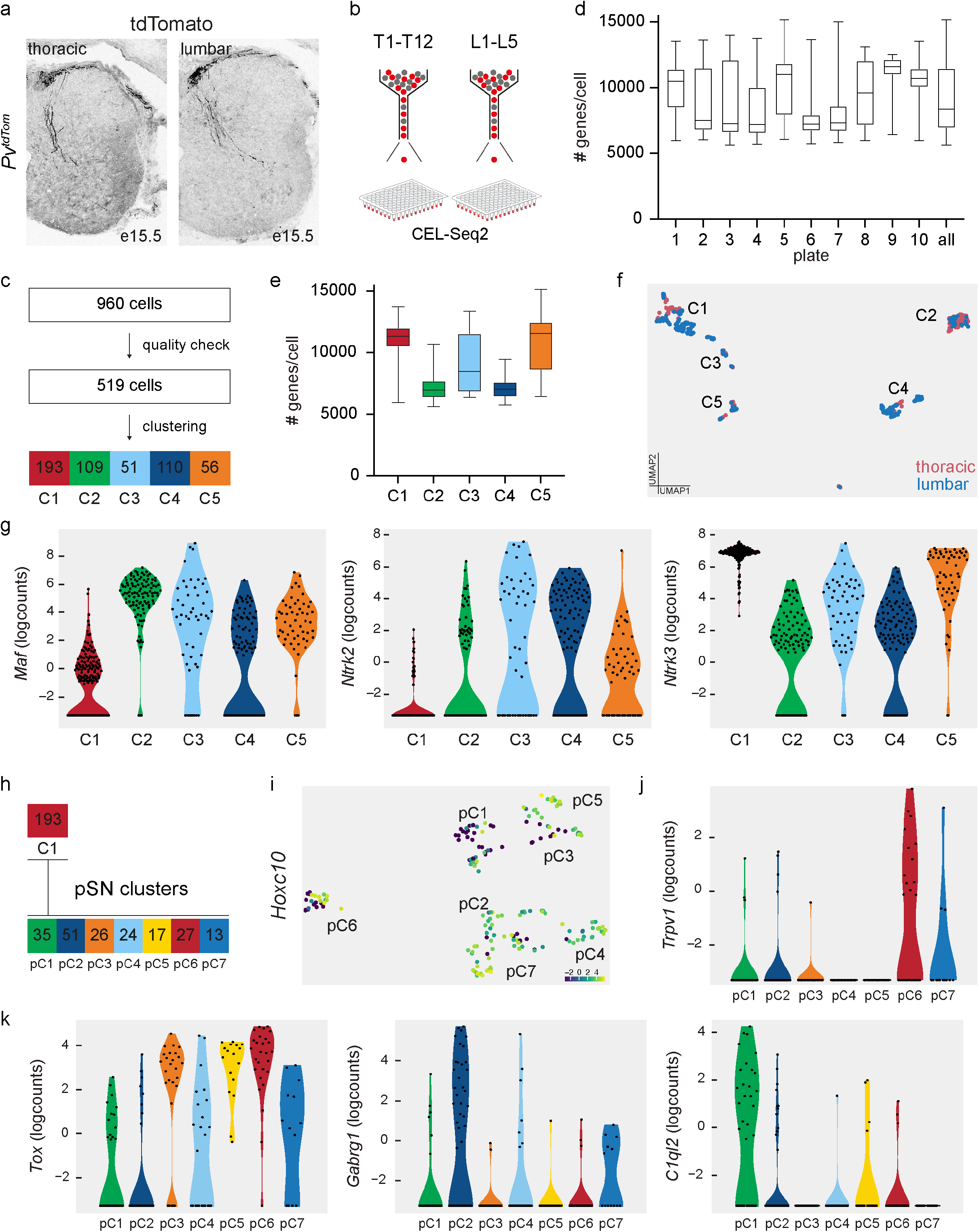
scRNA-seq analysis of e15.5 proprioceptors. **a)** Representative images of tdTomato^+^ afferents in thoracic and lumbar spinal cord from e15.5 *Pv^tdTom^* mice. **b)** Schematic representation of the single cell sorting strategy for sensory neurons dissociated from thoracic and lumbar DRG of e15.5 *Pv^tdTom^* mice. **c)** Number of e15.5 *Pv^tdTom^* DRG neurons sorted, analyzed after quality control and assigned to each cluster after bioinformatic analysis. **d)** Boxplots representing the number of genes per cells found in each 96 well plate (#1 to #10) and on average in all plates (“all”). **e)** Boxplots representing the number of genes per cells found in each cluster. **f)** UMAP visualization of proprioceptor clusters color coded according to thoracic (red) and lumbar (blue) origin of the cells. **g)** Violin plots showing expression (logcounts) of *Maf*, *Ntrk2,* and *Ntrk3* in clusters C1-C5. **h)** Number of C1 neurons re-clustered and assigned to proprioceptive clusters (pC) 1-7. **i)** UMAP visualization of *Hoxc10* expression (logcounts) in proprioceptor clusters. **j)** Violin plots showing expression (logcounts) of *Trpv1* in proprioceptor clusters pC1-pC7. **k)** Violin plots showing expression (logcounts) of *Tox, Gabrg1,* and *C1ql2* in proprioceptor clusters pC1-pC7.

**Extended Data Fig. 2.**
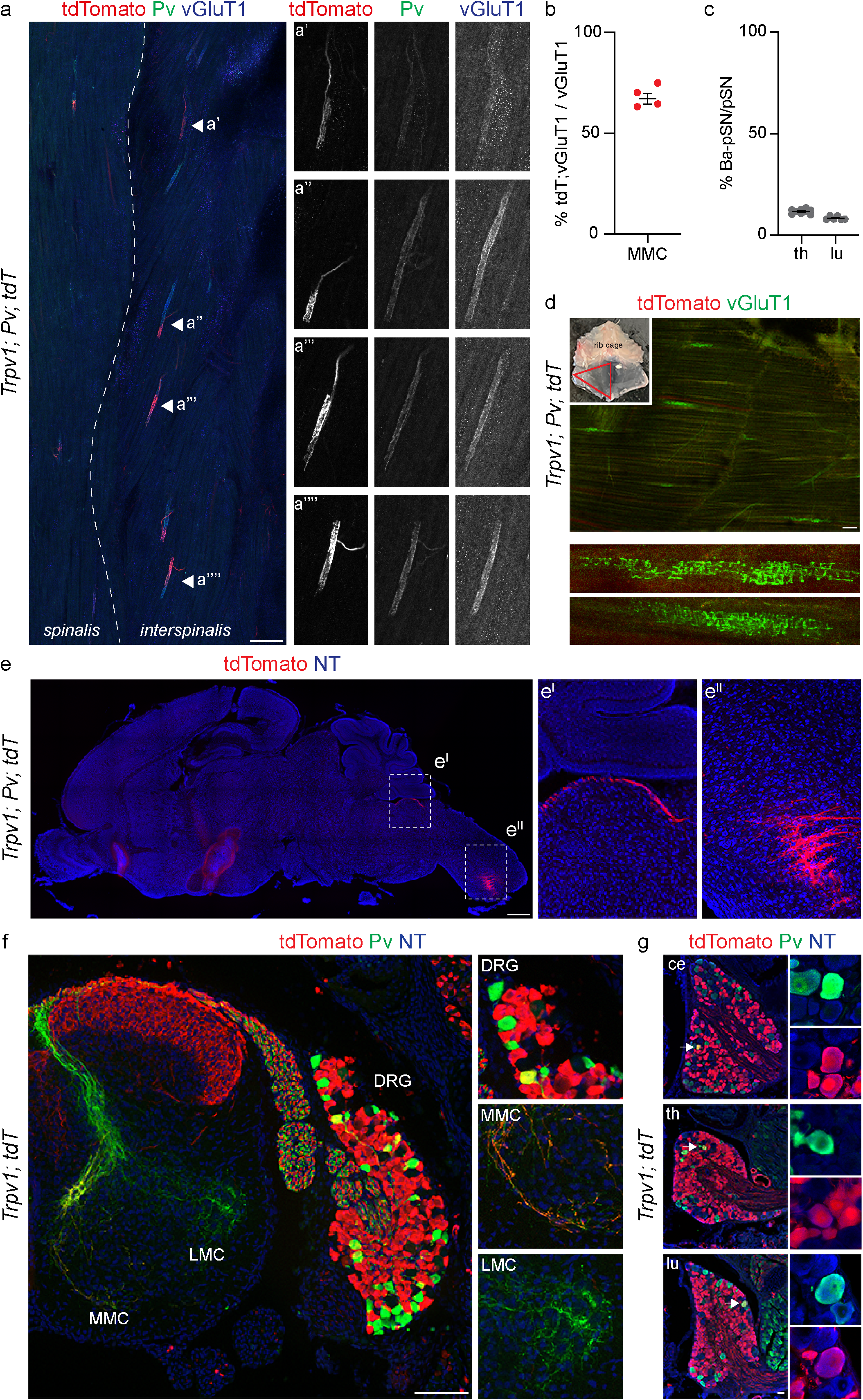
Genetic labelling of *Trpv1^+^* proprioceptors. **a)** Representative images of vGluT1^+^; Pv^+^; tdTomato^−^ spindles in the back (spinalis and interspinalis) muscles of p7 *Trpv1^Cre^; Pv^Flp^; Ai65* mice. Scale bar: 100 µm. **b)** Percentage of tdTomato^+^; vGluT1^+^ boutons juxtaposed to ChAT^+^ MMC neurons in p7 *Trpv1^Cre^; Pv^Flp^; Ai65* mice (n = 4 animals, 70 muscle spindles, mean ± SEM). **c)** Percentage of proprioceptors labelled in p7 *Trpv1^Cre^; Pv^Flp^; Ai65* mice at thoracic and lumbar levels. **d)** Representative image of vGluT1^+^; tdTomato^−^ spindles in the abdominal muscles of p7 *Trpv1^Cre^; Pv^Flp^; Ai65* mice. Scale bar: 100 µm. **e)** Representative sagittal brain section showing labelling of tdTomato^+^ proprioceptive afferents in the brainstem (e^I^) and cervical spinal cord (e^II^) of p7 *Trpv1^Cre^; Pv^Flp^; Ai65* mice. Scale bar: 500 µm. **f)** Representative image of sensory neurons labelled in a p7 *Trpv1^Cre^; Ai14* mice at lumbar level and high magnifications of DRG, MMC and LMC areas. Scale bar: 100 µm. **g)** Representative images of cervical (ce), thoracic (th), and lumbar (lu) DRG sections showing tdTomato^+^; Pv^+^ sensory neurons in p7 *Trpv1^Cre^; Ai14* mice. Scale bar: 25 µm.

**Extended Data Fig. 3.**
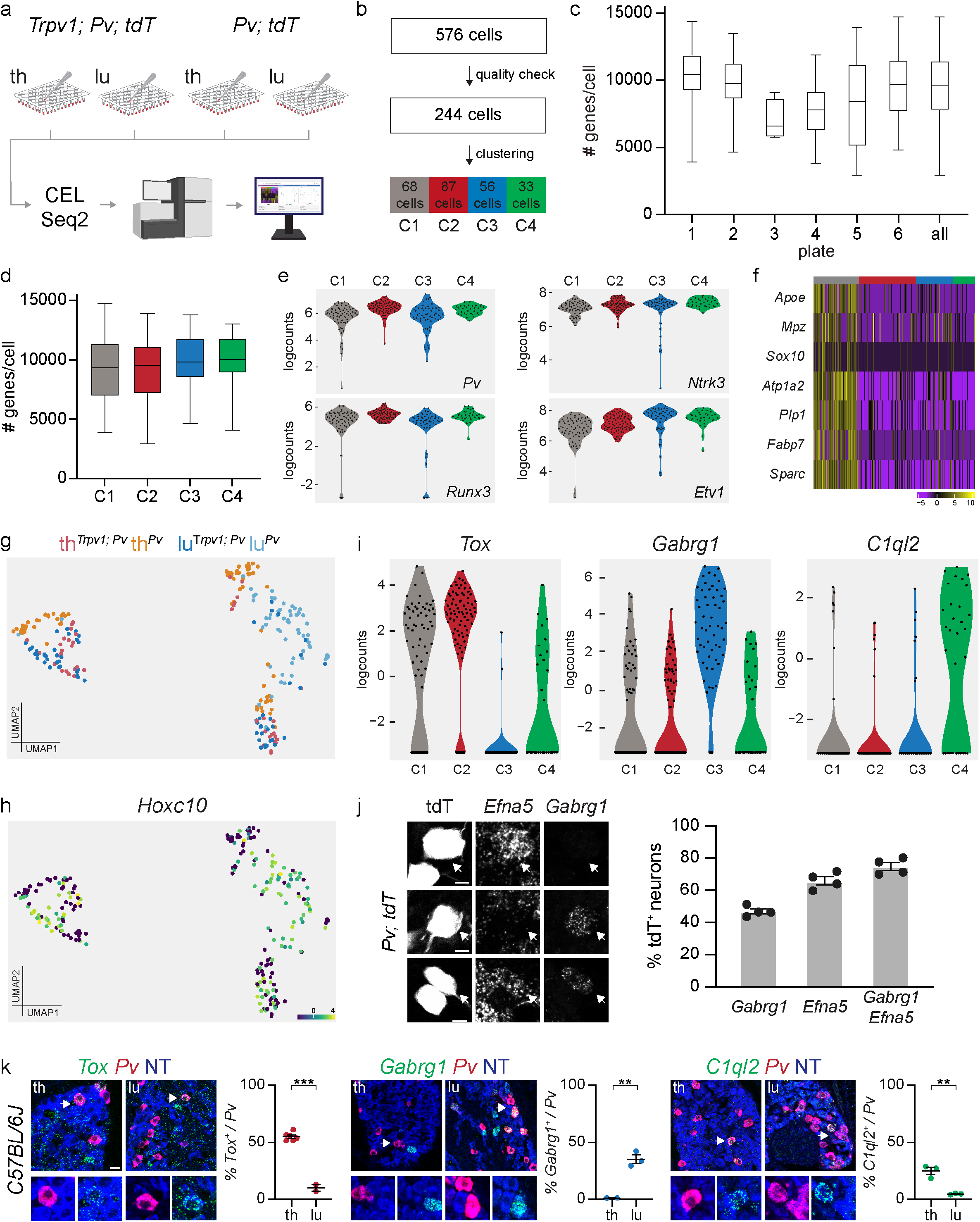
Transcriptome analysis at p1 and validation of proprioceptor muscle subtypes. **a)** Schematic representation of the single cell sorting strategy for neurons dissociated from p1 thoracic and lumbar DRG of *Trpv1^Cre^; Pv^Flp^; Ai65* and *Pv^Cre^; Ai14* mice. **b)** Number of tdTomato^+^ DRG neurons sorted, analyzed after quality control and assigned to each cluster after bioinformatic analysis. **c)** Boxplots representing the number of genes per cells found in each 96 well plate (#1 to #6) and on average (“all”). **d)** Boxplots representing the number of genes per cells found in each cluster. **e)** Violin plots showing expression (logcounts) of general proprioceptor markers (*Pv*, *Ntrk3*, *Runx3*, *Etv1*) at p1. **f)** Heatmap showing expression (logcounts) of glial cell markers in proprioceptor clusters at p1. **g)** UMAP visualization of clusters color coded to represent thoracic and lumbar origin of cells sorted from *Trpv1^Cre^; Pv^Flp^; Ai65* and *Pv^Cre^; Ai14* mice. **h)** UMAP visualization of *Hoxc10* expression (logcounts) in proprioceptor clusters at p1. **i)** Violin plots showing expression (logcounts) of *Tox*, *Gabrg,* and *C1ql2* in proprioceptor clusters at p1. **j)** Representative smFISH images of *Gabrg1* (C3) and *Efna5* (C3) expression in tdTomato^+^ lumbar DRG neurons of p1 *Pv^Cre^; Ai14* mice (left) and quantification of percentage of tdTomato^+^ sensory neurons expressing each marker alone or in combination (right). **k)** Representative smFISH images and quantification of *Tox* (C2), *Gabrg1* (C3), and *C1ql2* (C4) expression in Pv^+^ thoracic and lumbar DRG neurons of p1 wild type mice (each point represents one animal, mean ± SEM, T-test, ** p < 0.01, *** p < 0.001. Scale bar: 25 µm.

**Extended Data Fig. 4.**
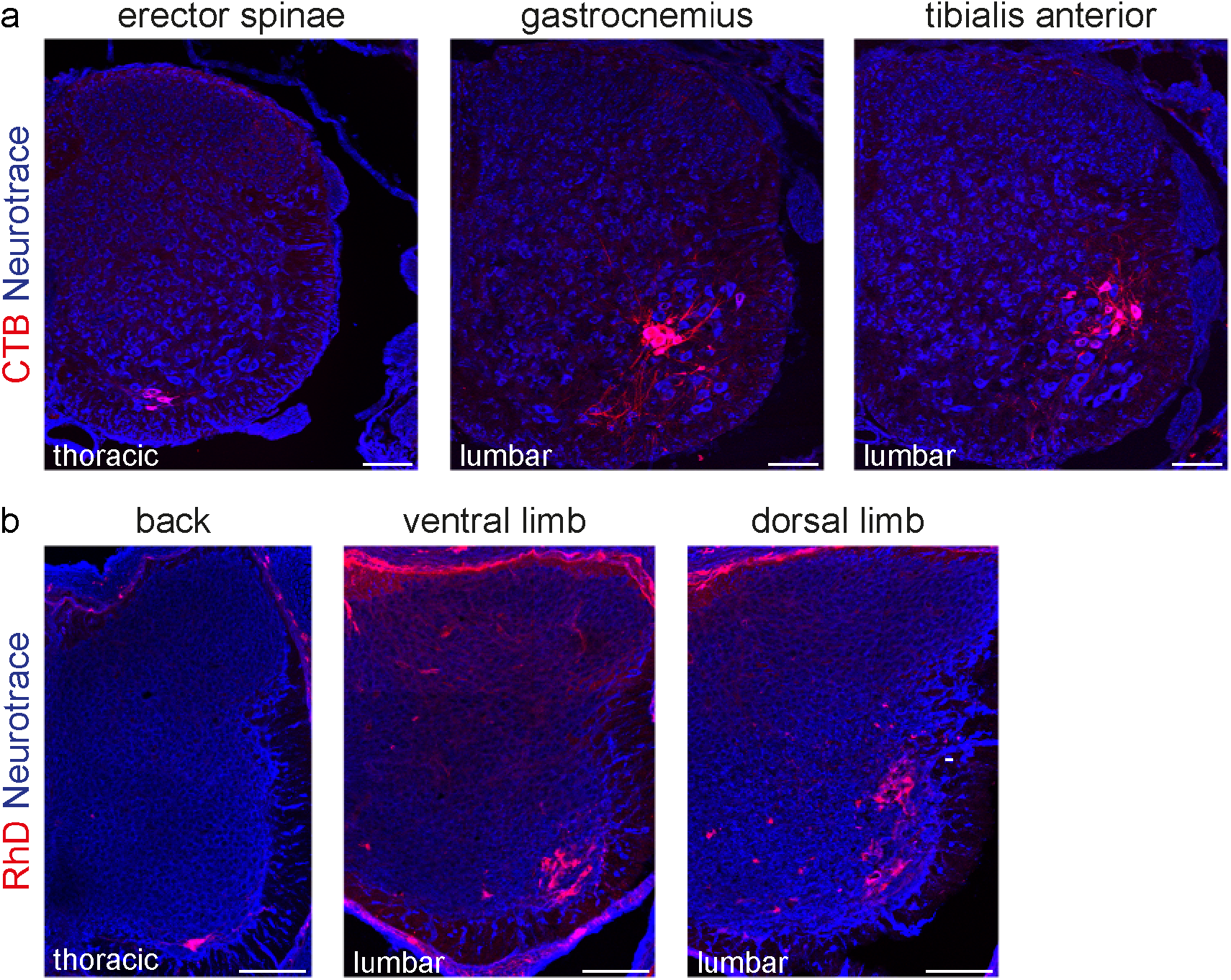
Retrograde labeling of epaxial and limb innervating motor neurons. **a)** Representative images of motor neurons retrogradely labelled after CTB injection in back (erector spinae) and hindlimb (gastrocnemius and tibialis anterior) muscles of wild-type mice. Scale bar:100 µm. **b)** Representative images of motor neurons retrogradely labelled after RhD injection in epaxial (back muscles) and ventral and dorsal hindlimb muscles of e15.5 wild-type embryos. Scale bar: 100 µm.

**Extended Data Fig. 5.**
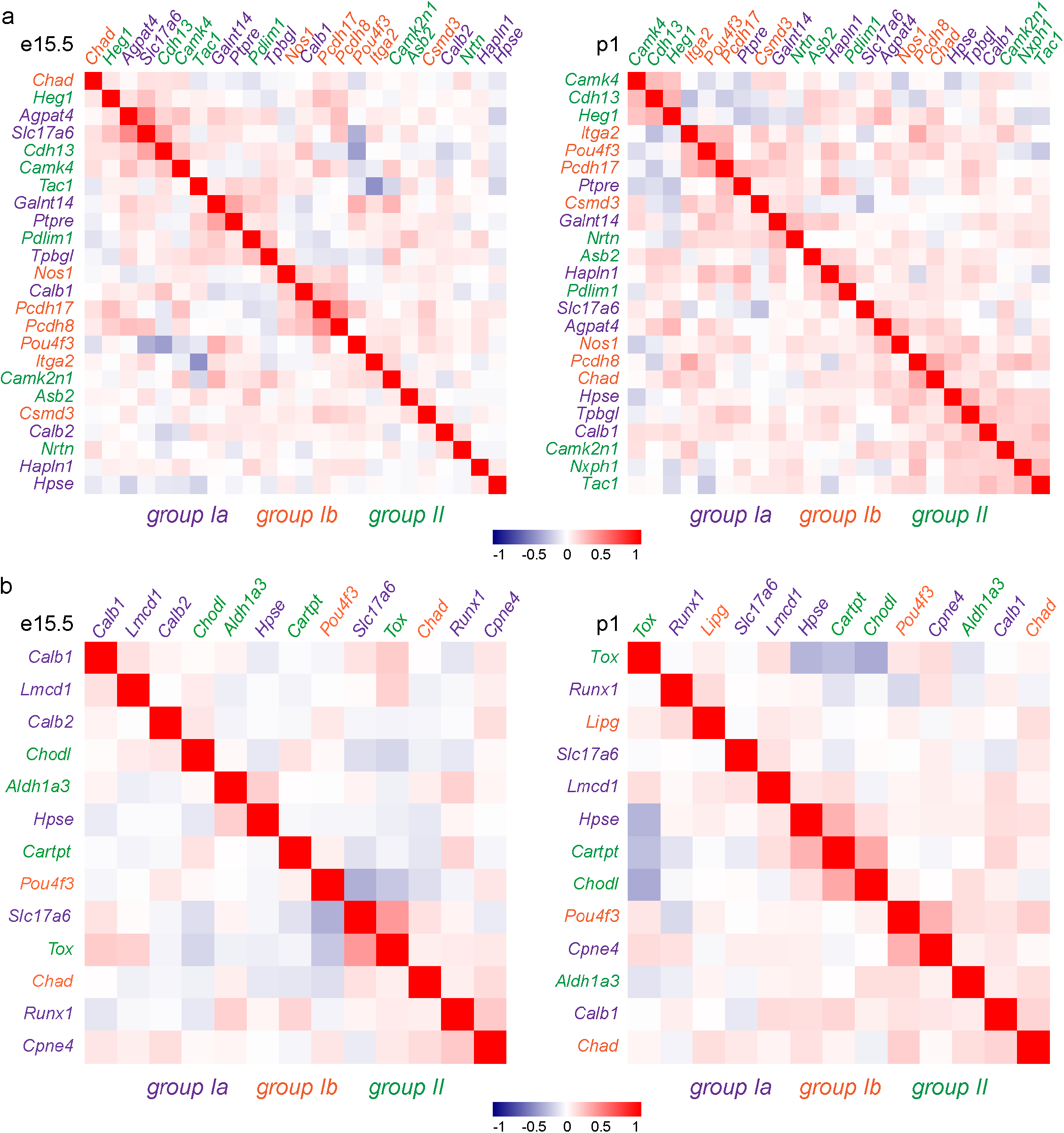
Signatures for “receptor-type” proprioceptors start emerging at p1. **a)** Heatmaps representing pairwise gene expression correlation values for group Ia (blue), group II (green), and group Ib (red) molecular signatures identified in *Oliver et al., 2021* at e15.5 (left) and p1 (right; Pearson’s r using logcounts). **b)** Heatmaps representing pairwise gene expression correlation values for group Ia (blue), group II (green), and group Ib (red) molecular signatures identified in *Wu et al., 2021* at e15.5 (left) and p1 (right; Pearson’s r using logcounts).

**Extended Data Fig. 6.**
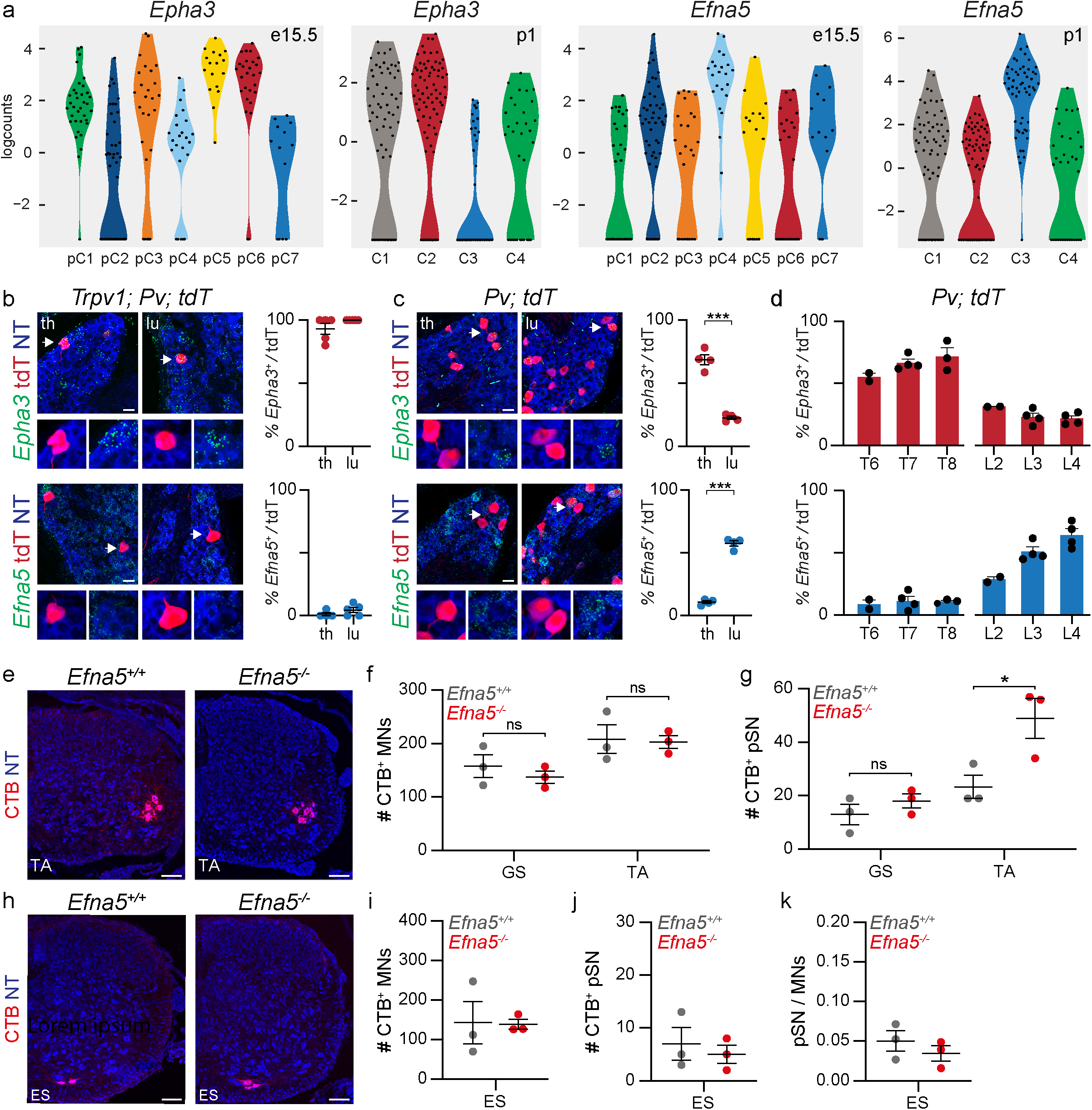
*Epha3* and *Efna5* expression defines proprioceptor muscle subtypes. **a)** Violin plots showing expression (logcounts) of *Epha3* and *Efna5* in proprioceptor clusters at e15.5 and p1. **b)** Representative smFISH images and quantification of *Epha3* (top, C2) and *Efna5* (bottom, C3) expression in tdTomato^+^ thoracic and lumbar DRG neurons of p1 *Trpv1^Cre^; Pv^Flp^; Ai65* mice (each point represents one animal, mean ± SEM). Scale bar: 25 µm. **c)** Representative smFISH images and quantification of *Epha3* (top, C2) and *Efna5* (bottom, C3) expression in tdTomato^+^ thoracic and lumbar DRG neurons of p1 *Pv^Cre^; Ai14* mice (each point represents one animal, mean ± SEM, t-test, *** p < 0.001). Scale bar: 25 µm. **d)** Distribution of *Epha3* (C2) and *Efna5* (C3) in tdTomato^+^ neurons from thoracic and lumbar DRG of p1 *Pv^Cre^; Ai14* mice (each point represents one animal, mean ± SEM). **e)** Representative images of motor neurons retrogradely labelled after CTB injection in the tibialis anterior (TA) muscle of p1 *Efna5^+/+^* and *Efna5^−/−^* mice. Scale bar: 100 µm. **f)** Number of motor neurons labelled after CTB injection in the gastrocnemius (GS) and tibialis anterior (TA) muscles of p1 *Efna5^+/+^* (gray) and *Efna5^−/−^* (red) mice (each point represents one animal, mean ± SEM, t-test, ns p > 0.05). **g)** Number of proprioceptors (*Pv^+^*) labelled after CTB injection in the gastrocnemius (GS) and tibialis anterior (TA) muscles of p1 *Efna5^+/+^* (gray) and *Efna5^−/−^* (red) mice (each point represents one animal, mean ± SEM, t-test, ns p > 0.05, * p < 0.05). **h)** Representative images of motor neurons retrogradely labelled after CTB injection in the erector spinae (ES) muscle of p1 *Efna5^+/+^* and *Efna5^−/−^* mice. Scale bar: 100 µm. **i)** Number of motor neurons labelled after CTB injection in the erector spinae (ES) muscle of p1 *Efna5^+/+^* (gray) and *Efna5^−/−^* (red) mice (each point represents one animal, mean ± SEM, t-test, ns p > 0.05). **j)** Number of proprioceptors (*Pv^+^*) labelled after CTB injection in the erector spinae (ES) muscle of p1 *Efna5^+/+^* (gray) and *Efna5^−/−^* (red) mice (each point represents one animal, mean ± SEM, t-test, ns p > 0.05). **k)** Ratio of proprioceptor (Pv^+^) per motor neuron labelled after CTB injection in the erector spinae (ES) muscle muscles of p1 *Efna5^+/+^* (gray) and *Efna5^−/−^* (red) mice (each point represents one animal, mean ± SEM, t-test, ns p > 0.05).

## Notes

### Competing Interest Statement

The authors have declared no competing interest.

